# Cycles of contamination and recovery: Combined sewer overflows drive acute but transient antimicrobial resistance exposure in an urban stream

**DOI:** 10.64898/2026.07.10.737760

**Authors:** Diala Konyali, Robin Pascal Mayer, Sara Schubert, David Kneis, Jakob Benisch, Geovanni Teran-Velasquez, Eda Deniz Erdem, Faina Tskhay, Reinhard Oertel, Peter Krebs, Thomas U. Berendonk, Uli Klümper

## Abstract

Combined sewer overflows (CSOs) are a major pathway for untreated wastewater into urban streams, yet their role in shaping antimicrobial resistance (AMR) dynamics remains poorly understood. Here, we used high-frequency, time-resolved sampling during two storm-triggered CSO events across two monitoring locations and one stormwater-only control site in an urban stream to quantify how these disturbances affect microbial communities, antibiotic resistance genes (ARGs), and mobile genetic elements (MGEs) in an urban stream.

CSO events caused rapid, up to two orders of magnitude, increases in bacterial, pathogen, and ARG abundance, with multiple transient peaks occurring within single overflow episodes. However, these increases were largely proportional to the total bacterial load, and most ARGs and MGEs did not change in relative abundance, indicating that CSOs primarily act as mass-transfer events rather than drivers of in situ selection. Downstream attenuation was governed by hydrological dilution despite additional CSO inputs: Both microbial and resistance signals largely returned to baseline within short time frames.

This demonstrates that CSOs function as hydrologically driven pulse disturbances that generate acute but transient AMR exposure. Because CSO events lack the sustained pressure associated with continuous wastewater discharges, rapid washout prevents the long-term establishment of sewage-derived resistance. These findings highlight that AMR risk in CSO-impacted systems is driven primarily by short-term exposure rather than by persistent ecological transformation, with important implications for urban water management under increasingly extreme rainfall conditions.

## 1. Introduction

Urban river ecosystems are shaped by natural hydrological processes and sustained anthropogenic pressures. Environmental surveillance of chemical and biological pollutants in these systems has therefore primarily focused on continuous point sources, particularly effluent from municipal wastewater treatment plants (WWTPs) [1,2]. Despite advanced treatment technologies, pathogens, antibiotic-resistant bacteria (ARB), and their resistance genes (ARGs) regularly reach and persist in receiving urban water bodies, reflecting the incomplete removal of microbial contaminants during wastewater treatment [3,4]. However, this focus on treated effluent overlooks other components of urban drainage systems that contribute intermittently but substantially to environmental contamination [5].

Increasing urbanization, combined with changing precipitation patterns under climate change, places additional pressure on ageing and often overloaded sewer infrastructure, even in regions with comparatively well-developed wastewater management [6–9]. One consequence of this is the more frequent activation of combined sewer overflows (CSOs), a designed feature of combined sewer networks that release diluted but untreated wastewater directly into receiving water bodies during heavy rainfall as a relief mechanism [5]. Even in temperate regions with generally high infrastructure standards and limited exposure to extreme weather, such as Dresden (Germany), CSO events still occur 15–20 times per year at major outlets [10]. This illustrates that episodic releases of untreated wastewater are a recurring, structurally embedded source of contamination, even under comparatively favourable conditions.

From an ecological perspective, combined sewer overflows can be understood as pulse disturbances, defined as sudden events that temporarily but strongly alter the physicochemical and biological conditions of receiving ecosystems [11,12]. During these events, urban streams are exposed to complex mixtures of contaminants, including potentially pathogenic and resistance-associated faecal bacteria, heavy metals, and pharmaceuticals that are usually present only at very low concentrations under baseline conditions [13–17]. The release of untreated wastewater during CSOs is therefore of particular concern because it bypasses the selective reduction processes of wastewater treatment, thereby directly introducing ARGs and human-associated pathogens into environmental microbial communities, where they may persist and interact with resident bacteria. Previous studies have suggested that the complex mixture of co-pollutants present in sewage may create conditions conducive to microbial stress and genetic exchange, potentially facilitating the spread of resistance determinants within receiving ecosystems [18,19]. As a result, CSOs may function not only as sources of resistant bacteria and genes but also as episodic disturbances that influence the dissemination of resistance beyond the immediate input event.

Although the effects of combined sewer overflows on general water quality parameters and their contribution to ARG loads have been confirmed, the temporal resolution of these impacts remains poorly understood [20–22]. Existing studies clearly identify CSOs as significant sources of resistance [17,23–25], yet it remains unclear how rapidly urban stream ecosystems recover following these disturbance events, or whether the apparent washout of contaminants masks longer-lasting alterations within the microbial community. Understanding recovery dynamics is critical, as repeated disturbance without full recovery could contribute to sustained alterations in environmental resistomes.

At the same time, the spatial impact of CSOs remains insufficiently understood. Urban streams often receive inputs from multiple discharge locations [10,12,26], generating overlapping contamination waves whose downstream effects depend on hydrological dilution, tributary inflows, and mixing processes. Disentangling the contributions of combined sewage from those of stormwater runoff is particularly important, as combined sewage introduces untreated wastewater that poses a substantially higher public health risk than stormwater runoff [5,27,28]. Quantifying how CSO-associated contamination attenuates or persists across spatial gradients is therefore essential for identifying high-risk zones and for informing effective monitoring and management strategies.

Thus, our goal in this study was to elucidate further how CSOs contribute to the spread and persistence of AMR in urbanized streams. To address the limited temporal and spatial resolution of previous studies, we investigated resistome dynamics under both seasonal baseline conditions and during high-frequency sampling campaigns triggered by the onset of storm-induced CSO events, using the Lockwitzbach stream in Dresden, Germany, as a model urban system. The Lockwitzbach flows approximately 24 km from rural headwaters into the urban area of Dresden, where increasing impervious surface cover coincides with multiple CSO discharge points [21]. By integrating the monitoring of two CSO-impacted river sections that differ in hydrological context, one located in the upper urban reach characterized by low baseflow and limited dilution capacity, and the other near the downstream end of the urban catchment where dilution is substantially higher, together with a stormwater-only control site, this system allowed us to assess both the immediate impacts of CSO events and the extent to which ARGs, pathogens, and mobile genetic elements (MGEs) attenuate or persist following disturbance. We provide a high-resolution characterization of how urban drainage infrastructure shapes environmental resistomes by combining 16S rRNA gene amplicon sequencing with high-throughput qPCR quantification of ARGs and MGEs and chemical analysis.

## 2. Materials and methods

### 2.1 Study Site

The study was conducted on Lockwitzbach, a 24 km urban tributary of the Elbe River in Dresden, Germany, with a catchment area of 87 km² [29]. The stream receives continuous effluent from the wastewater treatment plant in Kreischa (located ∼6 km upstream of the urban area) and, critically, intermittent discharges from an urban combined sewer network (Fig. S1). Two primary river monitoring stations were established: MS6 (50°59’00.1”N 13°48’02.1” E) captures the immediate impact of two CSO discharge points located approximately 660 m and 1 km upstream. MS4 (51°00’50.0”N 13°50’42.4” E) is located 6 km downstream, just before the confluence with the Elbe River, and integrates the cumulative inputs of an additional 12 CSO discharge points located in this stretch of the stream, with the sampling location 660 m downstream of the final discharge point. Distance to the last discharge point was selected so that sampling was representative of the state of the stream as high storm-induced turbulence and a distance of at least 660 m from the nearest CSO ensured sufficient mixing of the overflow plume prior to sampling (Fig. S1). In dry weather, water levels were comparable at both stations, averaging 10-13 cm. However, during active CSO events, the water levels rose sharply, with MS4 peaking substantially higher than MS6 (47 cm vs. 31 cm on 23 June). The stream width at MS4 is approximately double that at MS6 (3.6 m vs. 1.8 m). This is consistent with a large volume of water passing through MS4, establishing a clear dilution gradient.

### 2.2 Sample collection

Water samples were collected under two distinct conditions: seasonal baseline conditions during periods of normal flow with no recent rainfall or CSO activity, and CSO event-triggered sampling during heavy rainfall events. Under baseline conditions, samples were collected biweekly from January till September 2023. During overflow events, sampling was manually initiated at the onset of overflow conditions, identified by a rapid rise in water level, and the autosamplers then collected at fixed 15-minute intervals for the duration of the event. This study focuses on the two major rainfall events during the sampling period on 23. June and 12. July 2023, each delivering >15 mm of precipitation within few hours. During CSO events, automatic portable vacuum autosamplers were used at both stations (MAXX TP5 C at MS4; MAXX P6 at MS6, (MAXX, Rangendingen, Germany)), sampling directly from the stream, with intake hoses positioned approximately 0.5 m from the riverbank and co-located with the hydrological sensors. Recovery-phase samples were collected 2-3 days after each CSO event and were operationally designated as “recovery”. In total, 35 and 31 water samples were collected for MS6 and MS4, respectively, across all conditions and time points.

A third monitoring station (MS5, 51°00’17.9”N 13°50’45.2” E) was set up at inside a drain pipe discharging exclusively stormwater into the stream to differentiate between the contributions of combined sewer overflow and stormwater runoff to the resistome (Fig. S1). Located 4.5 km downstream of MS6, MS5 is not connected to the sanitary sewer system and only receives urban surface runoff during rainfall events. A total of 33 samples were collected at MS5 across rainfall events. By directly comparing the microbiome and resistome profiles of urban stormwater (MS5) with samples from the stream receiving combined sewage discharges (MS6, MS4), this design allowed source attribution of ARG inputs, distinguishing between faecal, wastewater-associated and predominantly environmental sources. Thus, MS5 functions as a source-attribution comparator rather than a temporal control: it indicates whether stormwater runoff alone can account for the observed ARG signals, but cannot by itself exclude other unmonitored inputs (Fig. S1).

### 2.3 Sample processing

Water samples were transported to the laboratory and vacuum-filtered through 0.22 μm polycarbonate membranes (47 mm, Sartorius, Göttingen, Germany) until clogging. The total filtered volume ranged from 300 to 400 mL, depending on turbidity. All filters were stored in sterile tubes at –20°C until DNA extraction. DNA extraction was performed using the DNeasy PowerSoil Pro Kit (Qiagen, Hilden, Germany). Filter membranes were aseptically cut, transferred to PowerBead tubes, and processed according to the manufacturer’s instructions. DNA concentration and purity (260/280 nm absorption ratio between 1.8–2.1) were determined using a NanoDrop 2000 spectrophotometer and DNA kept for downstream analysis at -20°C.

### 2.4 Quantification of chemical wastewater markers

Quantification of the targeted chemical markers, cotinine and metoprolol, was performed using solid-phase extraction (SPE) followed by liquid chromatography-tandem mass spectrometry (LC–MS/MS), based on methods described previously [30,31]. These compounds were chosen as anthropogenic indicators as cotinine was evaluated as a highly specific biomarker of human domestic wastewater (derived from nicotine consumption) and metoprolol was evaluated as a representative pharmaceutical beta-blocker [32–35].

Homogenized water samples (50 mL) were dosed with Na₂EDTA (0.8 mg mL⁻¹). Following shaking, centrifugation, and filtration through glass-fibre filters (<0.7 µm; WICOM, Heppenheim, Germany), the filtrates were adjusted to pH 3.5±0.2 with formic acid (LC–MS grade; Sigma, St. Louis, MO, USA). For calibration, a blank matrix (diluted and adjusted to pH 3.5±0.2) was spiked with isotope-labelled internal standards and native analytes. Sample extraction was carried out using Oasis HLB SPE cartridges (30 mg, Waters, Milford, MA, USA) on a Gilson ASPEC GX-271 automated SPE system (Middleton, WI, USA). Reconstituted dried extracts were analysed using an LC–MS/MS system equipped with a Kinetex® RP column (2.6 µm, 150 × 3.0 mm) and a C18 SecurityGuard cartridge (4 × 2 mm; both Phenomenex, Aschaffenburg, Germany). Chromatographic separation was achieved under reversed-phase conditions. Detection was performed on an API 4000 triple quadrupole mass spectrometer (AB Sciex, Framingham, MA, USA) equipped with an electrospray ionisation (ESI) source operating in multiple reaction monitoring (MRM) mode. Quantification relied on external calibration with isotope-labelled internal standards. Method quantification limits were 10 ng L⁻¹ for both cotinine and metoprolol. Acceptance criteria for the analyses included a signal-to-noise ratio greater than 10 and intra- and inter-day precision of less than 20% deviation.

### 2.5 Absolute bacterial abundance

For absolute bacterial abundance, qPCR targeting the 16S rRNA gene was performed using the primers F: 5′-TCCTACGGGAGGCAGCAGT-3′, R: 5′-ATTACCGCGGCTGCTGG-3′ [36].

Reactions (20 μL) contained 10 μL of Luna Universal qPCR Master Mix (New England Biolabs) with SYBR Green detection, 5 μL of template DNA (10 ng total), and were run in triplicate on a C1000 Touch Thermal Cycler (Bio-Rad USA). The qPCR protocol consisted of 95°C for 10 min followed by 40 cycles of 95°C for 15 s and 60°C for 1 min. Standard curves were generated using linearized pNORM plasmid DNA [37]. Only reactions with amplification efficiency between 90% and 110% and standard curves with R² ≥ 0.99 were accepted.

### 2.6 High-throughput qPCR for ARG and MGE quantification

To quantify the relative abundance of ARGs and MGEs, DNA samples were analysed at Resistomap Oy (Helsinki, Finland) using a SmartChip real-time PCR system. The protocol included primer sets targeting 24 ARGs, 3 MGE marker genes, and the 16S rRNA gene (Table S1). In addition, three targets were also selected to provide species and genus level resolution of pathogenic bacteria. *Escherichia coli* and *Enterococcus spp.* were included as canonical faecal indicator bacteria regulated under WHO guidelines for freshwater systems [38]. *Klebsiella pneumoniae* was selected as a representative WHO critical priority opportunistic pathogen [39], as its carbapenem-resistant lineages are of clinical concern in wastewater-impacted aquatic environments [40]. Amplification was performed at 95°C for 10 min, followed by 40 cycles of 95°C for 30 s and 60°C for 30 s. The detection limit was set at Ct 27 [41], and the limit of quantification was set at 25 gene copies per reaction. Relative abundance was normalized to the 16S rRNA gene using the ΔCt method [42]. ARGs were chosen for clinical relevance, human health risk, and consistent detection in wastewater, spanning major resistance classes (β-lactams including carbapenemases and ESBLs, macrolides, tetracyclines, sulfonamides, quinolones, aminoglycosides, trimethoprim, vancomycin, and colistin); the full panel and target sequences are listed in Table S1.

### 2.7 16S rRNA Gene Amplicon Sequencing

To analyse the bacterial community composition, 16S rRNA gene amplicon sequencing targeting the V3–V4 region (F: 5′-CCTACGGGAGGCAGCAG-3′; R: 5′-GGACTACHVGGGTWTCTAAT-3′) [43] was conducted on the Illumina NovaSeq platform at the IKMB sequencing facility (Kiel University, Germany) with a target depth of ≥10,000 reads per sample. The 16S rRNA sequences were processed using Mothur (v1.48.0) following the MiSeq SOP [44,45]. Briefly, raw reads were merged. Low-quality and chimeric sequences were removed, and the remaining reads were aligned against the SILVA 138 reference database [46,47] and classified using the RDP database [48]. Eukaryotic, chloroplast, archaeal, and mitochondrial sequences were excluded.

Operational Taxonomic Units (OTUs) were defined at 97% similarity and rarefied to the minimum read depth to allow comparison of microbial diversity and taxonomic composition across samples.

### 2.8 Pathogen classification

To determine the percentage of potential bacterial pathogens, an ASV-based rather than OTU-based analysis was performed to achieve higher accuracy. Sequences were processed in QIIME2 (v2023.9) [49]. Briefly, demultiplexing and primer trimming (cutadapt) [50] were performed, then ASVs were generated by denoising using DADA2 [51]. A pretrained Naïve Bayes classifier trained on the SILVA 138 database was used for taxonomic annotation. The percentage of potential pathogens was then calculated by comparing the identified ASVs against a reference list of 1,513 human-pathogenic bacteria [52].

### 2.9 Statistical analysis

Data visualization and statistical analysis were conducted in R (version 4.3.1; R Development Core Team). Given the pronounced right-skew typical of environmental microbial abundance data, non-parametric tests were used throughout. The Mann-Whitney U test was used to compare total bacterial, pathogen, ARG, and MGE abundances between seasonal baseline and CSO conditions; this test is equivalent to the Wilcoxon rank-sum test, and for brevity it is referred to as the Mann-Whitney U test throughout. For comparisons across three hydrological phases (e.g., pre-CSO baseline, CSO peak, and recovery), the Kruskal-Wallis rank-sum test was applied, followed by Dunn’s post hoc test. Crucially, to account for multiple hypothesis testing across the variety of monitored ARGs and MGEs, p-values were adjusted using the Benjamini-Hochberg False Discovery Rate (FDR) procedure. Adjusted p-values<0.05 were considered statistically significant. Box plots display medians and interquartile ranges; group means are reported in the text to support fold-change calculations. Fold changes were calculated as the ratio of group means (mean CSO abundance divided by mean seasonal baseline abundance, where the seasonal baseline comprises all seasonal samples) to allow direct comparison across all measured analytes. For the season-versus-CSO comparisons, all seasonal samples were pooled to represent the range of unperturbed baseline conditions across the study period. For the three-phase recovery analysis, the pre-event seasonal sampling (season1) was used as the pre-event baseline to isolate the immediate pre-CSO state from post-event recovery samples.

Non-metric Multidimensional Scaling (NMDS) was performed with the Vegan package [53]. Euclidean distance was used for ARG analysis and Bray-Curtis dissimilarity for OTUs. ANOSIM was used to determine the significant differences between groups. To identify bacterial taxa significantly associated with CSO events versus seasonal conditions, Linear Discriminant Analysis Effect Size (LEfSe) was performed separately for MS6 and MS4 using the microbiomeMarker package (v1.7.0) in R [54]. OTU tables and taxonomies were imported into phyloseq objects [55] for each monitoring site. The LEfSe algorithm applies a three-step approach to identify differentially abundant features. First, the Kruskal-Wallis rank-sum test identifies features with significant differences in abundance between conditions (α=0.05). Pairwise Wilcoxon rank-sum tests confirm biological consistency (α=0.05), and finally, Linear Discriminant Analysis estimates the effect size of each differentially abundant taxon [56]. Only taxa with an LDA score ≥ 3.0 were considered biologically relevant, a conventional threshold corresponding to a strong and consistent discriminatory effect between conditions. Multiple testing correction was applied using the Benjamini-Hochberg false discovery rate method. LEfSe was performed across all taxonomic levels, from phylum to genus; unclassified or uncultured taxa were excluded from the final results to focus on biologically interpretable lineages.

## 3. Results

To investigate how CSO events shape the urban resistome, we compared microbial community composition, antibiotic resistance genes (ARGs), and mobile genetic elements (MGEs) between seasonal baseline samples and samples collected during two storm-triggered CSO events at three monitoring stations: MS6 (an upstream CSO-impacted site), MS4 (a downstream site located 6 km below MS6, immediately before the confluence with the Elbe River), and MS5 (stormwater-only discharge point).

### 3.1 CSO events are defined by sharp hydrological surges

Under baseline dry-weather conditions, water levels were similar at both monitoring stations, and sampling was done every two weeks. In contrast, CSO event sampling at 15-minute intervals was manually initiated during active overflow. Two precipitation events exceeding 15 mm rainfall within 24 hours occurred on 23 June and 12 July 2023, triggering CSO discharges in the urban catchment during the study period. During these events, water levels at MS4 reached 47 cm (23 June) and 29 cm (12 July), compared to 31 cm and 14 cm at MS6, respectively (Fig. S2).

### 3.2 Chemical markers confirm untreated wastewater influx during CSOs

To independently verify that the introduction of untreated sewage drove the hydrological surges during CSO events, we quantified two highly specific anthropogenic chemical markers: the human metabolite cotinine and the pharmaceutical metoprolol [32–35]. Under seasonal baseline conditions, concentrations remained relatively low. During active CSO events, however, concentrations spiked substantially. Cotinine significantly increased 5-fold from a baseline of 6.09±7.6 ng/L to 30.12±13.46 ng/L during overflows (P=0.029, Mann–Whitney U test), while metoprolol significantly increased 2.2-fold from 31.34±9.56 ng/L to 68.70±37.63 ng/L (P=0.035). The enrichment was even more pronounced at the downstream station MS4, where cotinine significantly increased 18-fold (baseline: 6.01±5.02 ng/L; CSO: 111.25±69.51 ng/L; P=0.019) and metoprolol increased 4.7-fold (baseline: 31.82±31.37 ng/L; CSO: 150.8±50.77 ng/L; P=0.032). The consistent increase of both human-specific markers across both sites provides clear, non-biological confirmation that the storm-triggered CSOs heavily contaminate the river with raw wastewater.

### 3.3 CSO-driven increases in bacterial and potential pathogen abundance

To determine how strongly CSO events alter the stream’s microbial composition, we quantified total bacterial and potential bacterial pathogen abundance under baseline and CSO conditions at both monitoring stations. Quantification of 16S rRNA gene copies revealed a pronounced increase in total bacterial abundance during CSO events. However, during each CSO event, bacterial abundance did not follow a simple rise-and-fall pattern (Fig. 1A, B). Instead, high-frequency sampling revealed multiple transient peaks within a single overflow episode, with concentrations increasing and decreasing repeatedly over the course of the event. These intra-event oscillations likely reflect changes in sewer discharge and variable contributions of combined sewage during different intensities of rainfall. Thus, CSO events act not as single, uniform pulses but as multiple short-lived disturbances.

**Figure 1:**
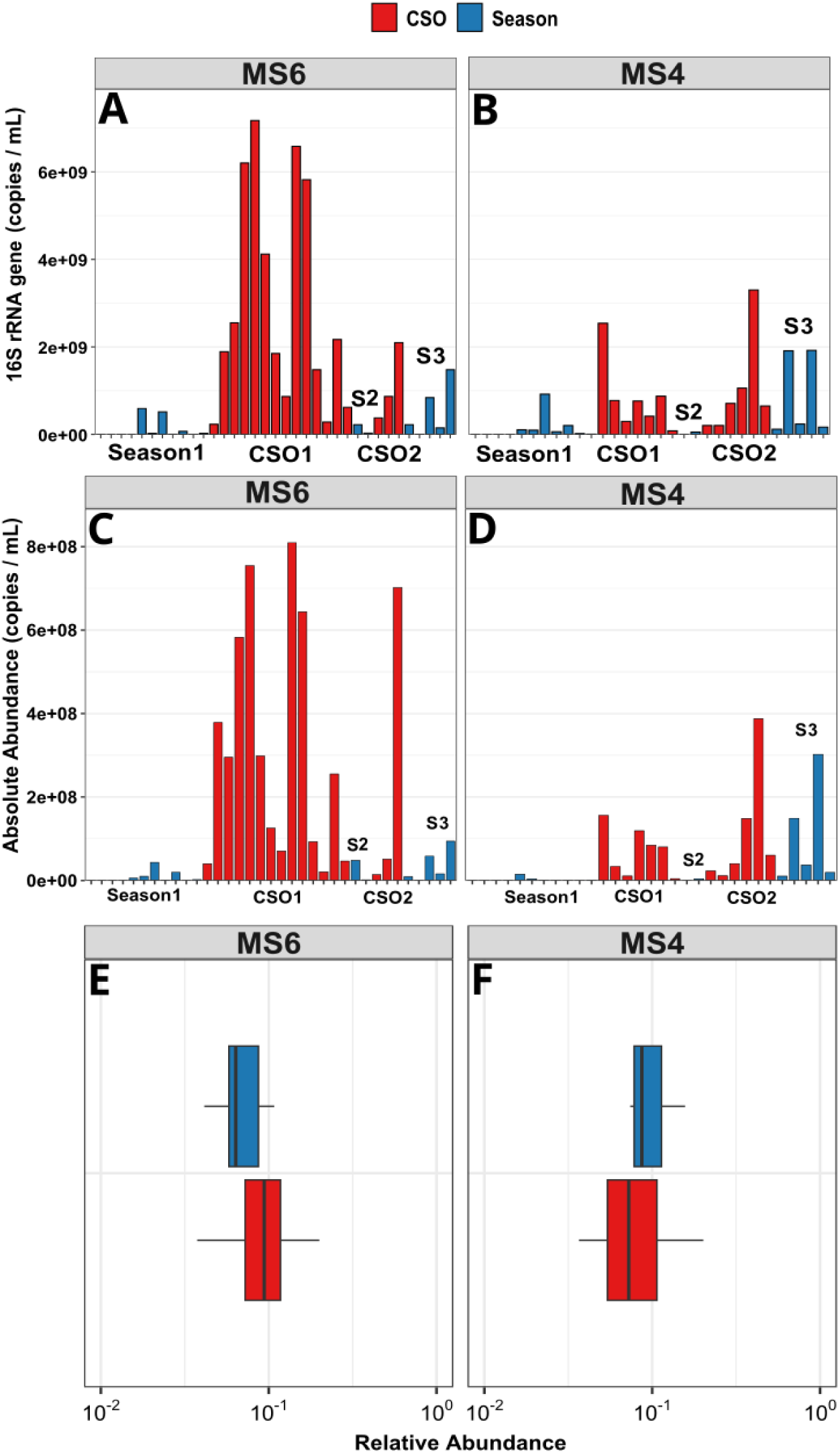
Total bacterial and potential pathogen abundances during seasonal baseline and CSO conditions. (A, B) Total bacterial abundance (16S rRNA gene copies/mL) quantified by qPCR at the upstream station MS6 (A) and downstream station MS4 (B). (C, D) Absolute abundance of potential human pathogens (copies/mL), based on classification against 1,513 clinically relevant bacterial pathogens, at MS6 (C) and MS4 (D). (E, F) Relative abundance of potential pathogens, expressed as a ratio of the total bacterial community, at MS6 (E) and MS4 (F). Box plots display the median (centre line), interquartile range (box), and 1.5× interquartile range (whiskers). Across all panels, seasonal baseline conditions are denoted in blue and CSO events in red.

At the upstream site MS6, bacterial abundance increased on average 11-fold during CSOs (2.66±2.38 × 10^9^ copies/mL) compared to baseline conditions (2.33±3.96 × 10^8^ copies/mL; Mann–Whitney U test, P=6.62 × 10^-6^, nseason=18, nCSO=17) (Fig. 1A), reaching concentrations typical of raw wastewater (10^9^–10^10^ copies/mL). At MS4, bacterial abundance increased on average 3-fold during CSO events (9.15±9.50 × 10^8^ copies/mL) relative to baseline (3.24±6.16 × 10^8^ copies/mL; P=2.19 × 10^-3^, nseason=18, nCSO=13) (Fig. 1B). Although substantial, this enrichment was markedly lower than at MS6, consistent with hydrological dilution and increased flow along the 6 km stretch between the two sites. Despite the larger absolute rise in water level at MS4, the wider channel (3.6 m vs. 1.8 m at MS6) and with MS4 being located downstream of additional CSO outlets, the dilution effect dominates, moderating contamination inputs at this location (Fig. 1B).

Potential pathogen abundance followed the same intra-event oscillatory and spatial patterns. At MS6, absolute abundance of potential pathogens increased 24-fold during CSO events (3.05±2.87 × 10^8^ copies/mL) compared to baseline (1.29±1.85 × 10^7^ copies/mL; Mann–Whitney U test, P=5.24 × 10^-6^, nseason=18, nCSO=17) (Fig. 1C). At MS4, potential pathogen abundance increased 3-fold (3.01±7.64 × 10^7^ vs. 8.93±10.34 × 10^7^ copies/mL; P=1.03 × 10^-3^, nseason=18, nCSO=13) (Fig. 1D), confirming that CSO events introduce substantial potential pathogen loads throughout the urban area. However, no clear dilution effect was visible here.

Despite these pronounced absolute increases, the proportion of potential human-pathogenic genera within the bacterial community remained stable at both sites. At MS6, the relative abundance of potential human-pathogenic genera did not differ significantly between baseline and CSO conditions (10.56±9.22% vs. 11.20±7.05%; Mann–Whitney U test, P=0.382, Fig. 1E), which was consistent with MS4 (6.93±6.13% vs. 9.38±4.98%; P=0.307; Fig. 1F). The apparent increase at MS4 did not reach significance, and the modest sample size limits the statistical power to detect a proportional shift of this magnitude. These findings indicate that the incoming untreated sewage possesses a similar proportion of potential human-pathogenic genera as the stream, which is already influenced by continuous wastewater effluent release approximately 7 km upstream of MS6. While the relative abundance remained stable, the absolute abundance of potential pathogens increased by up to 24-fold. Because the chemical markers were quantified only at the event level rather than at the 15-minute biological sampling interval, we do not infer peak-to-peak temporal alignment; nonetheless, the concurrent enrichment of human-specific markers (cotinine and metoprolol) during the same events indicates that the observed surges are driven by the introduction of wastewater rather than the mere resuspension of sediment-associated microbiota.

To complement the amplicon-based pathogen classification, which relies on genus-level resolution, we targeted three specific pathogen markers included in the SmartChip qPCR panel: *E. coli, Enterococci,* and *K. pneumoniae.* All three increased significantly in absolute abundance during CSO events at both stations (Mann–Whitney U test, FDR-BH; adjusted P<0.05 for all; Fig. S3). In terms of relative abundance, *E. coli* increased significantly at both MS6 (adjusted P=0.005) and MS4 (adjusted P=0.039), and *Enterococci* at MS4 (adjusted P=0.039), indicating that these organisms were disproportionately concentrated in the CSO inputs relative to the resident stream community. *K. pneumoniae* showed no significant relative change at either site (adjusted P>0.05). These targeted findings confirm and extend the broad pathogen enrichment observed through amplicon-based classification, providing species-level evidence that CSO events introduce primary faecal indicators and a WHO critical-priority pathogen into the receiving stream.

Together, these results demonstrate that CSO events are associated with rapid increases in total microbial and pathogen abundance. Despite MS4 being located downstream of additional CSO outlets, the magnitude of enrichment for bacteria was consistently lower at MS4 than at MS6, consistent with downstream dilution. This suggests that the increased water level reaching MS4 attenuates the CSO effect. The observed increases in bacterial abundance provide a quantitative basis for subsequent analysis of ARG abundance and broader resistome dynamics.

### 3.3 CSO-driven effects on the river resistome

#### 3.3.1 Absolute ARG abundance

The absolute abundance of ARGs was quantified across sites and conditions to determine if the resistome mirrored the sharp increases in the bacterial community. Absolute ARG abundance displayed intra-event variability similar to bacterial abundance, with multiple peaks observed within a single overflow event (Fig. 2A, B). On average, total ARG abundance increased significantly during CSO events at both monitoring stations. At MS6, total ARG abundance increased 11-fold during CSOs relative to all seasonal samples (Season: 1.03 × 10^8^±1.56 × 10^8^ copies/mL; CSO: 1.16 × 10^9^±1.02 × 10^9^ copies/mL; Mann–Whitney U test, P=2.98 × 10^-5^, nseason=18, nCSO=17; Fig. 2A). At MS4, a significant but more modest 2-fold increase was observed (Season: 2.38 × 10^8^±4.69 × 10^8^ copies/mL vs. CSO: 4.44 × 10^8^±5.72 × 10^8^ copies/mL; P=0.029, nseason=18, nCSO=13; Fig. 2B). This spatial gradient mirrors the pattern observed for total bacterial abundance and is consistent with downstream dilution of sewage-derived inputs.

**Figure 2:**
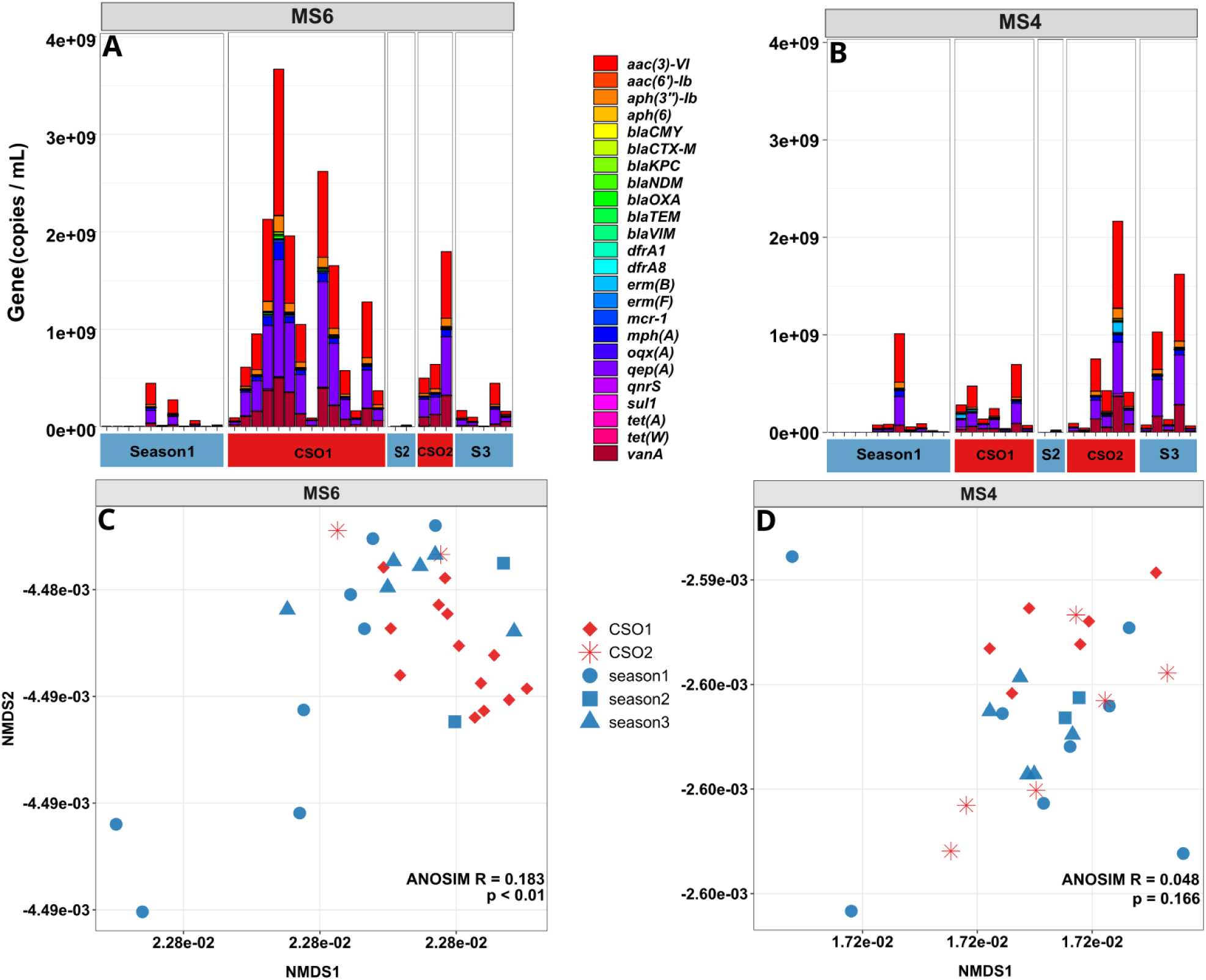
Absolute abundance and beta-diversity of ARGs at monitoring stations MS6 and MS4. (A, B) Absolute ARG abundance at the upstream station MS6 (A) and downstream station MS4 (B) under seasonal baseline and CSO conditions. (C, D) Non-metric multidimensional scaling (NMDS) ordination based on Euclidean distances of ARG abundance profiles at MS6 (C) and MS4 (D), illustrating the shifts in resistome composition between hydrological phases.

At MS6 all 24 quantified ARGs were significantly enriched during CSO events (Mann–Whitney U tests with Benjamini–Hochberg FDR correction; adjusted P<0.05 for all genes, Fig. 3A). *erm*B exhibited the strongest enrichment at 34-fold, followed by *aac(6’)-*Ib (21-fold), *van*A (20-fold), *bla*OXA (19-fold), and *sul*1 (16-fold). The lowest enrichment was *bla*KPC at 5-fold. Together, these results confirm a broad and pronounced surge in ARG concentration across all resistance classes at the upstream site, paralleling the increase in total bacterial abundance.

**Figure 3:**
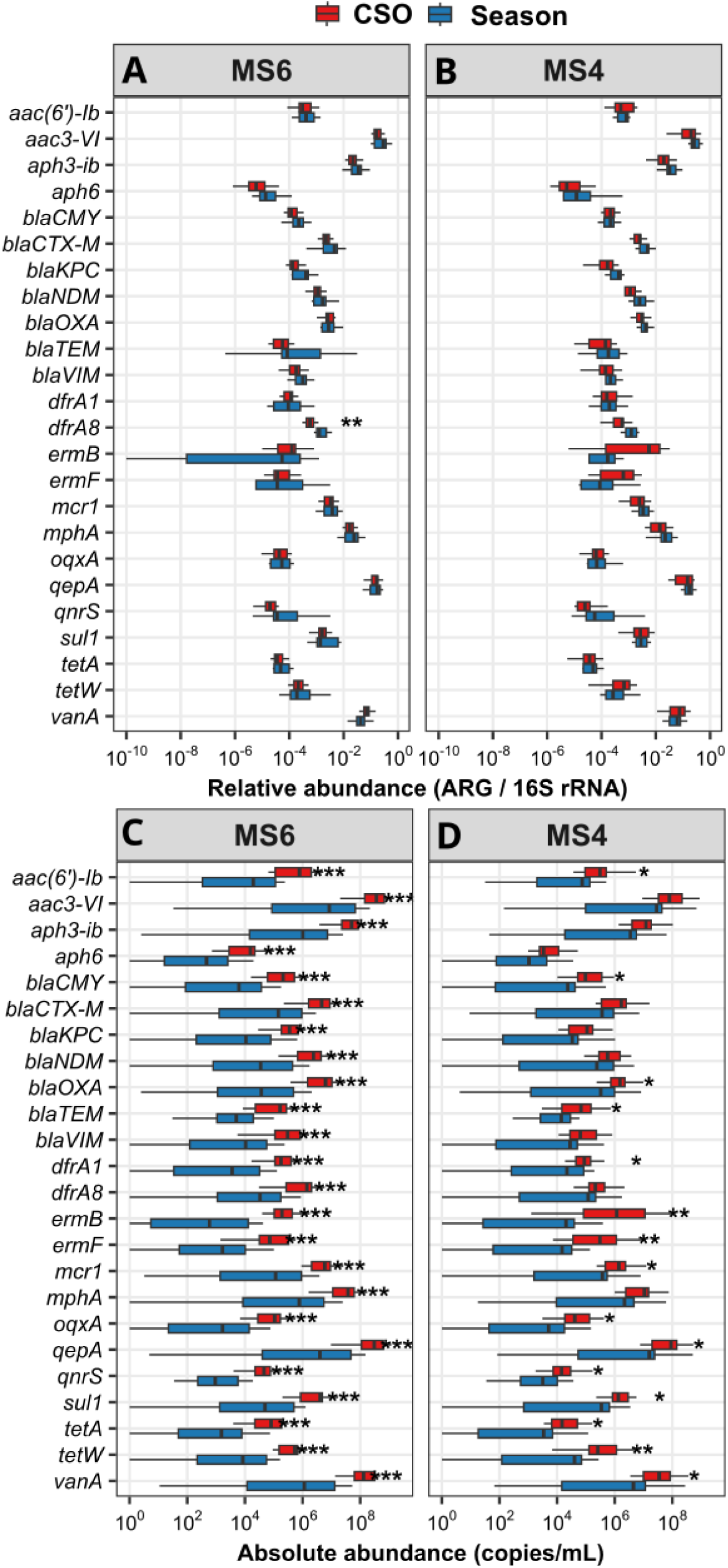
Absolute and relative abundance of ARGs during seasonal baseline and CSO conditions. (A, B) Absolute ARG abundance at MS6 (A) and MS4 (B). (C, D) Relative ARG abundance at the upstream station MS6 (C) and downstream station MS4 (D). Across all panels, seasonal baseline conditions are denoted in blue and CSO events in red. Box plots display the median (center line), interquartile range (box), and 1.5× interquartile range (whiskers). Asterisks indicate statistical significance after Benjamini–Hochberg FDR correction (*adjusted P<0.05, **adjusted P<0.01, ***adjusted P <0.001).

Overall, 15 of 24 ARGs were significantly enriched during CSO events at MS4 (adjusted P<0.05; Fig. 3B). *erm*B again showed the highest fold enrichment (241-fold; from a baseline mean of 6.2 × 10^4^ to 1.5 × 10^7^ copies/mL), followed by *erm*F (56-fold). The exceptionally large fold change for *erm*B reflects its very low and frequently non-detectable abundance in baseline samples at this site, which inflates the ratio of group means; consistent with this, *erm*B showed no significant change in relative abundance after FDR correction (Fig. 3D). Consistent with the high dilution at MS4, the remaining significant ARGs displayed more modest fold changes of 2–16-fold. Nine ARGs did not reach statistical significance: *bla*NDM, *bla*KPC, *bla*VIM, *bla*CTX-M, *dfr*A8, *aac3-*VI, *mph*A, *aph3-*ib, and *aph*6 (adjusted P>0.05 for all). The combination of reduced absolute concentrations of these genes and the high downstream dilution likely lowered their signal below statistical detectability. Nevertheless, significant resistome contamination during overflow conditions was confirmed by the overall 1.9-fold increase in total ARG load at MS4.

#### 3.3.2 Relative ARG abundance

Whereas absolute ARG abundance increased uniformly during CSO events, analysis of relative abundance revealed that most genes remained proportionally stable, consistent with the pattern observed for pathogen relative abundance. Most ARGs, including broadly distributed ARGs such as *sul*1 and *tet*W, displayed no significant differences in relative abundance between CSO and seasonal baseline samples (Mann–Whitney U test; Fig. 3C, D).

At the upstream station MS6, only the trimethoprim resistance gene *dfr*A8 showed a significant decrease in relative abundance during CSO events (3.9-fold; adjusted P=0.007; Fig. 3C). This suggests that, compared to the incoming combined sewage, the resident stream community was proportionately more enriched for *dfr*A8. This supports the interpretation that CSOs act primarily as mass-transfer events rather than drivers of in-stream resistance selection.

At the downstream station MS4, no ARG showed a significant change in relative abundance after FDR correction (adjusted P>0.05 for all comparisons; Fig. 3D). Together with the MS6 results, the near-complete stability of relative ARG abundance at both sites confirms that the surges in absolute ARG concentrations observed during CSOs reflect the dilution of stream water with incoming sewage rather than selective amplification of specific resistance determinants within the receiving community.

#### 3.3.3 Resistome composition shifts during CSO events

To evaluate whether CSO events altered the overall resistome structure, we performed NMDS using Euclidean distances among ARG abundance profiles. At MS6, samples collected during CSO events showed separation from baseline samples, and ANOSIM confirmed a significant difference in resistome composition (R=0.162, P=0.005; Fig. 2C). Although the effect size was moderate, this separation indicates that CSO discharges measurably altered ARG composition at the upstream site. In contrast, no significant difference was detected at MS4 (R=0.021, P=0.278; Fig. 2D), suggesting that downstream dilution reduced microbial composition shifts despite substantial increases in total ARG load.

#### 3.3.4 CSO events, rather than stormwater runoff, drive ARG enrichment

To further clarify the origin of this ARG enrichment, and because river samples collected during CSO events receive a mixture of combined sewage and stormwater runoff, we next evaluated whether stormwater influx alone could explain the observed increases in ARG. For this, we compared ARG abundance in stormwater-only discharge (MS5) with both CSO-impacted samples (MS6 and MS4 during overflow events) and seasonal baseline conditions.

ARG abundances at MS5 were significantly lower than those measured during CSO events at both MS6 (23 of 24 genes, adjusted P<0.05) and MS4 (19 of 24 genes, adjusted P<0.01; Mann–Whitney U test with FDR correction) (Fig. 4). In contrast, when compared to seasonal baseline samples at MS6 and MS4, the MS5 resistome showed strong similarity. No significant differences were detected for 23 of 24 ARGs. The macrolide resistance gene *mph*A was the only exception, occurring at significantly lower abundance in MS5 relative to baseline (adjusted P<1 × 10^-5^). This indicates that stormwater runoff alone does not account for the observed ARG surges during CSO events. This conclusion is supported by recent physicochemical profiling of stormwater outlets in this specific Dresden catchment, which demonstrated that stormwater runoff is predominantly composed of fine inorganic sediments and trace metals, rather than biological or organic loads [57]. Instead, combined sewage inputs represent the most likely source of ARG enrichment in the urban stream.

**Figure 4:**
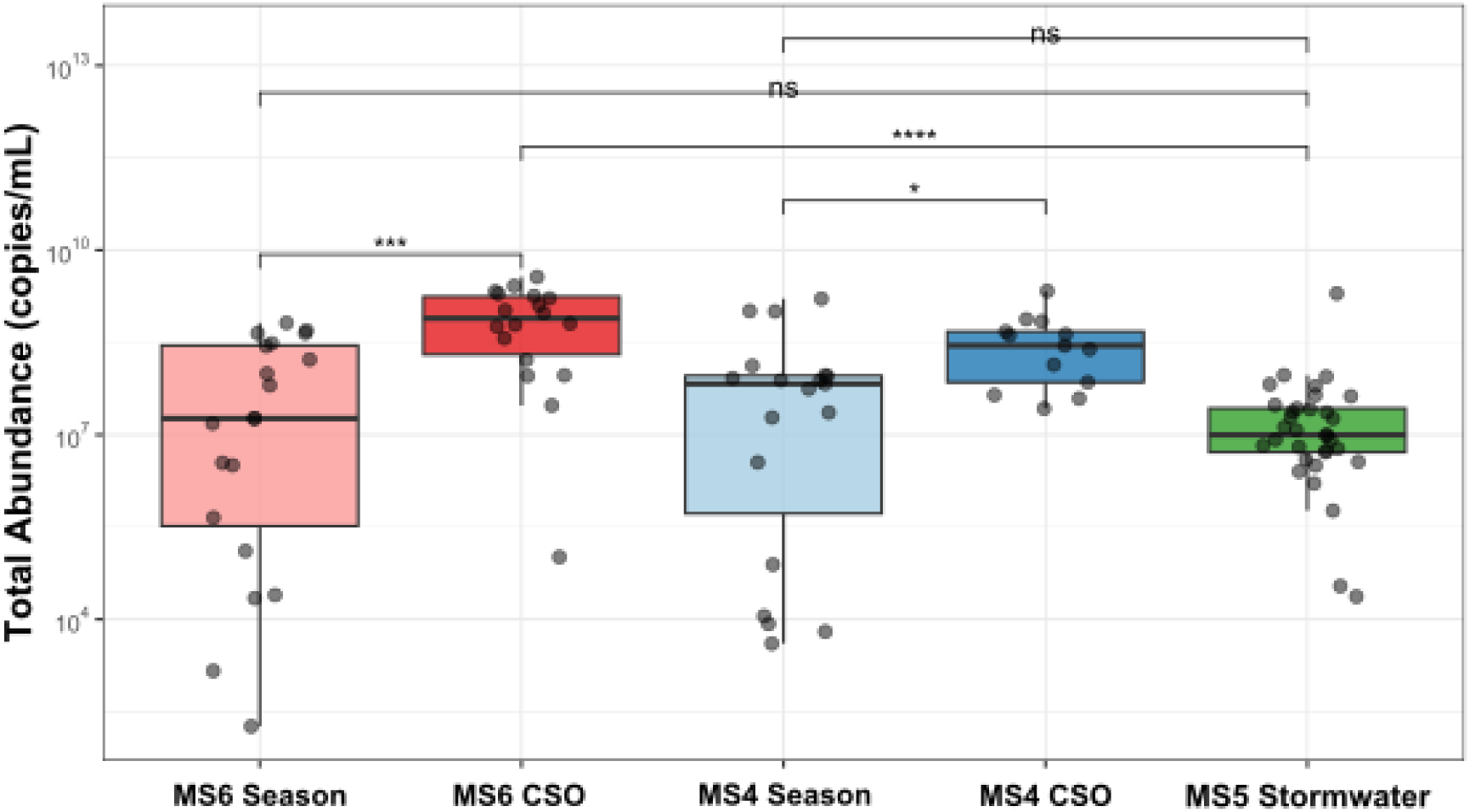
Comparison of ARG abundance between stormwater-only discharge (MS5, green), CSO-impacted sites (MS6 and MS4, red, blue), and seasonal baseline conditions (pink, light blue) for total ARG abundance. **adjusted P<0.01, ***adjusted P<0.001 (Mann-Whitney U test with FDR correction).

### 3.4 CSO events transiently elevate plasmid-associated mobility markers

Because the public health risk associated with ARG enrichment depends not only on gene abundance but also on their capacity for horizontal transfer [58], we next quantified three broad-host-range plasmid incompatibility groups (IncP, IncW, and IncQ) that mediate HGT. Absolute abundance of IncP plasmids increased significantly by approximately 10-fold at MS6 (adjusted P<0.001; Fig. 5) and 1.6-fold at MS4 (adjusted P=0.030) during CSO events compared to baseline conditions. Similarly, IncW plasmids increased by 7-fold at MS6 and 2.1-fold at MS4 (adjusted P<0.001 and 0.032, respectively), following the same upstream-to-downstream attenuation pattern observed for total ARG load. IncQ plasmids were likewise significantly enriched, increasing by 7-fold at MS6 and 3.0-fold at MS4 during overflow conditions (adjusted P<0.001 and adjusted P=0.021, respectively). Most plasmid comparisons showed no significant change in relative abundance. Only IncQ decreased slightly but significantly at MS6 (adjusted P=0.0001), similar to IncW at MS6 (adjusted P=0.039). The significant relative decreases in IncQ and IncW abundance at MS6, alongside the otherwise stable relative abundances, collectively indicate that absolute plasmid peaks during CSO events primarily reflect a mass influx of plasmid-carrying bacteria rather than selective amplification within the resident community.

**Figure 5:**
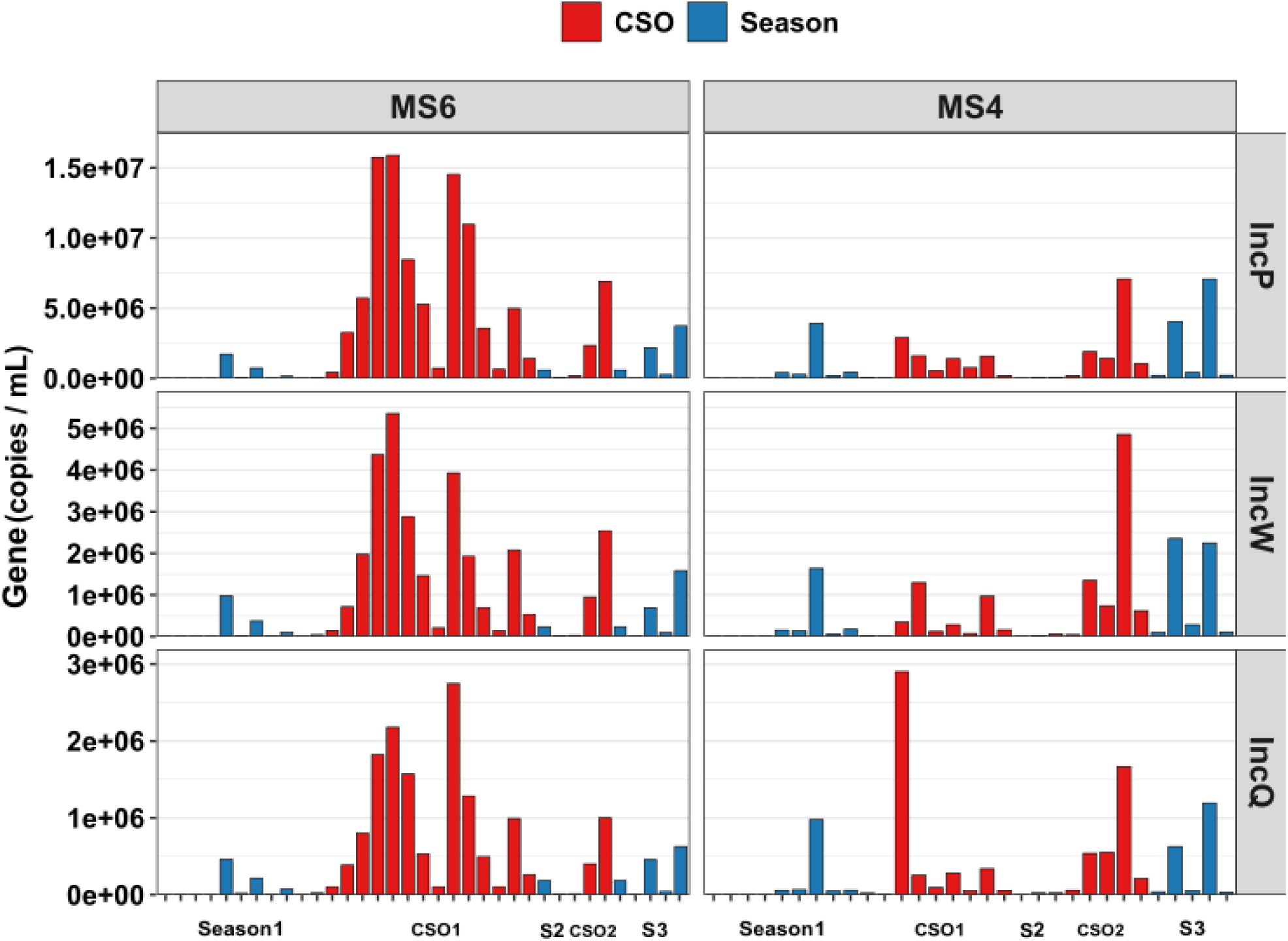
Abundance of MGEs during seasonal baseline and CSO conditions. Bar graphs displaying the abundance of plasmid replicons (A) IncP, (B) IncW, and (C) IncQ at the upstream (MS6) and downstream (MS4) monitoring stations. Across all panels, seasonal baseline conditions are denoted in blue and CSO events in red.

### 3.5 CSO events induce limited shifts in microbial community structure

Despite the substantial influx of sewage-associated bacteria during CSO events, overall community structure remained comparatively stable. NMDS based on Bray-Curtis dissimilarities revealed only modest differences between baseline and CSO samples (Fig. 6A, B). At MS6, CSO samples formed a weakly distinct cluster from seasonal samples (ANOSIM R=0.102, P=0.026), indicating a statistically significant but ecologically small shift. At MS4, no significant separation was observed (R=-0.04, P=0.782), reflecting near-complete overlap. The larger effect size at MS6 than at MS4 mirrors the gradient in bacterial enrichment and suggests partial attenuation of disturbance downstream. Consistent with this structural resilience, phylum-level community composition remained stable during CSO events (Fig. 6C, D). Under baseline conditions, communities at both sites were dominated by *Pseudomonadota* (MS6: 36.4 ± 19.8%, MS4: 47.8±20.1%), *Acidobacteriota* (MS6: 21.7 ±11.9%, MS4: 17.3±18.6%), *Actinomycetota* (MS6: 16.5 ± 12%, MS4:11.3± 12%), and *Planctomycetota (MS6: 1.1±2.3%, MS4: 6.7±*12.9%). During CSO events, the relative proportions of these dominant phyla did not differ significantly from those under seasonal conditions at either site (P>0.05 for all comparisons), indicating that sewage inputs did not disrupt the broad taxonomic structure. To identify taxa specifically associated with CSO events versus baseline conditions, Linear Discriminant Analysis Effect Size (LEfSe) was performed at both monitoring sites. At MS6, 36 taxa showed significant differential abundance between CSO and seasonal conditions (LDA>3.0, adjusted p<0.05), with 27 taxa enriched during CSO events and 9 enriched under seasonal conditions. At the downstream site MS4, only 18 taxa were differentially abundant (12 CSO-enriched, 6 seasonally-enriched), reflecting the dilution-driven attenuation of CSO impacts observed in bacterial abundance and ARG loads.

**Figure 6:**
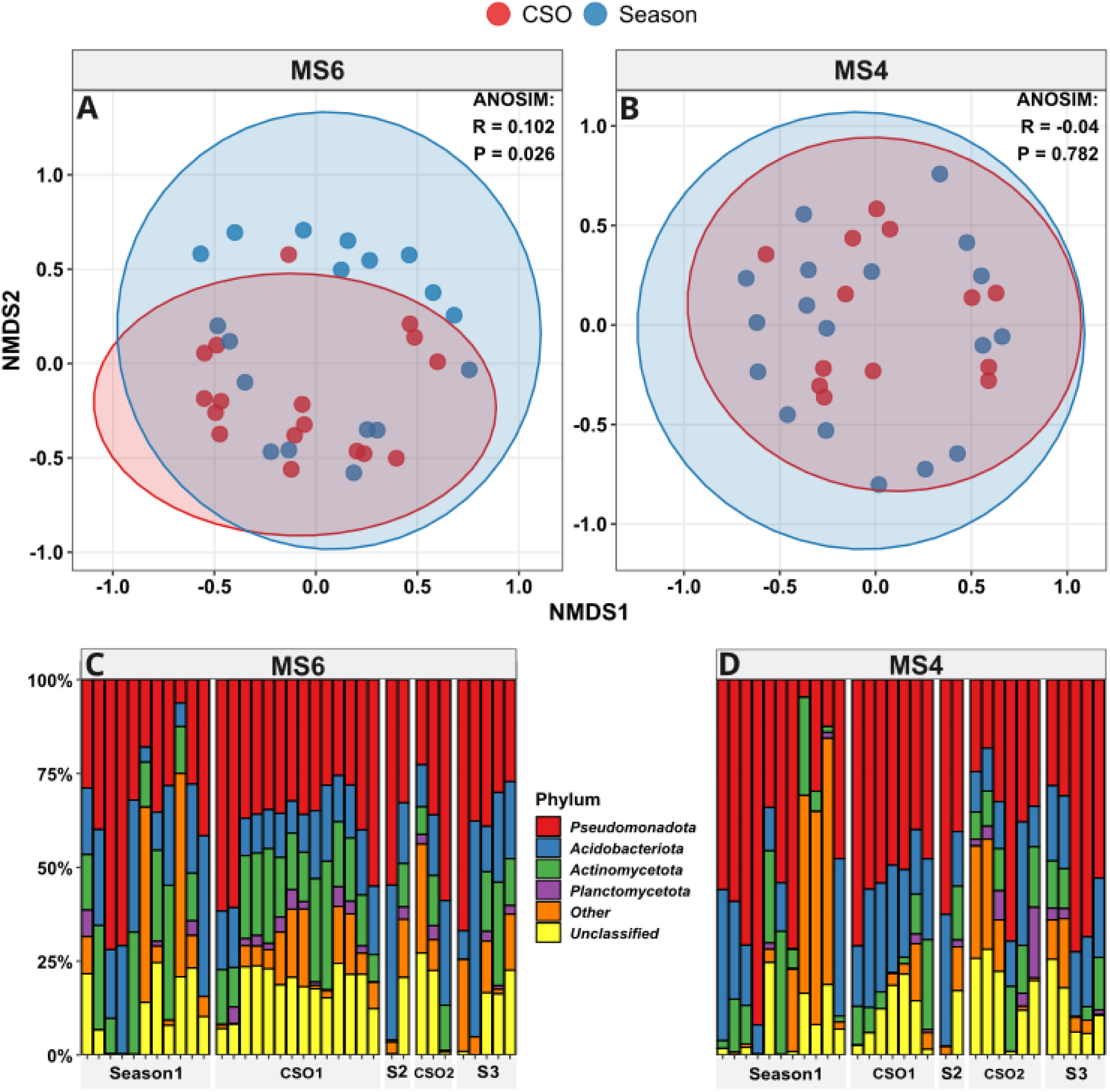
Bacterial community structure and taxonomic composition during seasonal baseline and CSO conditions. (A, B) Non-metric multidimensional scaling (NMDS) ordination of Bray-Curtis dissimilarities based on operational taxonomic unit (OTU) relative abundance at the upstream station MS6 (A) and downstream station MS4 (B). Seasonal baseline samples are represented by blue dots and CSO event samples by red dots, with ellipses denoting 95% confidence intervals. ANOSIM statistics are provided to indicate the significance of community dissimilarity at each site. (C, D) Phylum-level taxonomic composition illustrating the relative abundance (percentage) of dominant bacterial phyla at MS6 (C) and MS4 (D).

Importantly, despite these statistically significant shifts in specific lineages, the overall microbial community structure was consistent, as evidenced by the small effect sizes and the absence of phylum-level compositional changes (Fig. S4). The LEfSe results are consistent with the transient introduction of sewage-associated taxa during CSO events, rather than with fundamental restructuring of the stream microbiome.

### 3.6 ARG and pathogen abundance return toward baseline after CSO events

To assess the duration of CSO impacts and the resilience of the stream ecosystem, we compared ARG and pathogen abundance across three phases: pre-CSO baseline, CSO peak, and post-CSO recovery. At MS6, total ARG abundance during recovery (1.48±1.71 × 10^8^ copies/mL, n=7) declined significantly from CSO peak levels (Dunn’s post-hoc, P=0.015) and did not differ significantly from the pre-event baseline (P=0.386), indicating complete normalisation following overflow cessation. At the gene level, all 24 ARGs increased significantly from baseline to CSO peak (adjusted P<0.05). During recovery, 21 of 24 ARGs declined significantly from CSO peak levels (adjusted P<0.05 for these 21); the three remaining genes (*aph*6, *bla*TEM, *erm*F) had already returned close to baseline, so the CSO-vs-recovery contrast was not significant for them. Critically, no ARG differed significantly from the pre-event baseline during recovery (adjusted P>0.05 for all 24 genes; Fig. 7A), confirming that the resistome returned to its pre-disturbance composition. Pathogen abundance at MS6 followed the same pattern, declining significantly from CSO levels during recovery (P=0.012) and returning to baseline (P=0.231; Fig. S5A). Notably, even during the overflow phase itself, concentrations repeatedly declined toward near-baseline levels between transient intra-event peaks (Fig. 1A-D), indicating rapid washout over short hydrological periods. The consistent post-event return to baseline reflects the same dilution-driven dynamics observed during the oscillatory phase of the CSO event.

**Figure 7:**
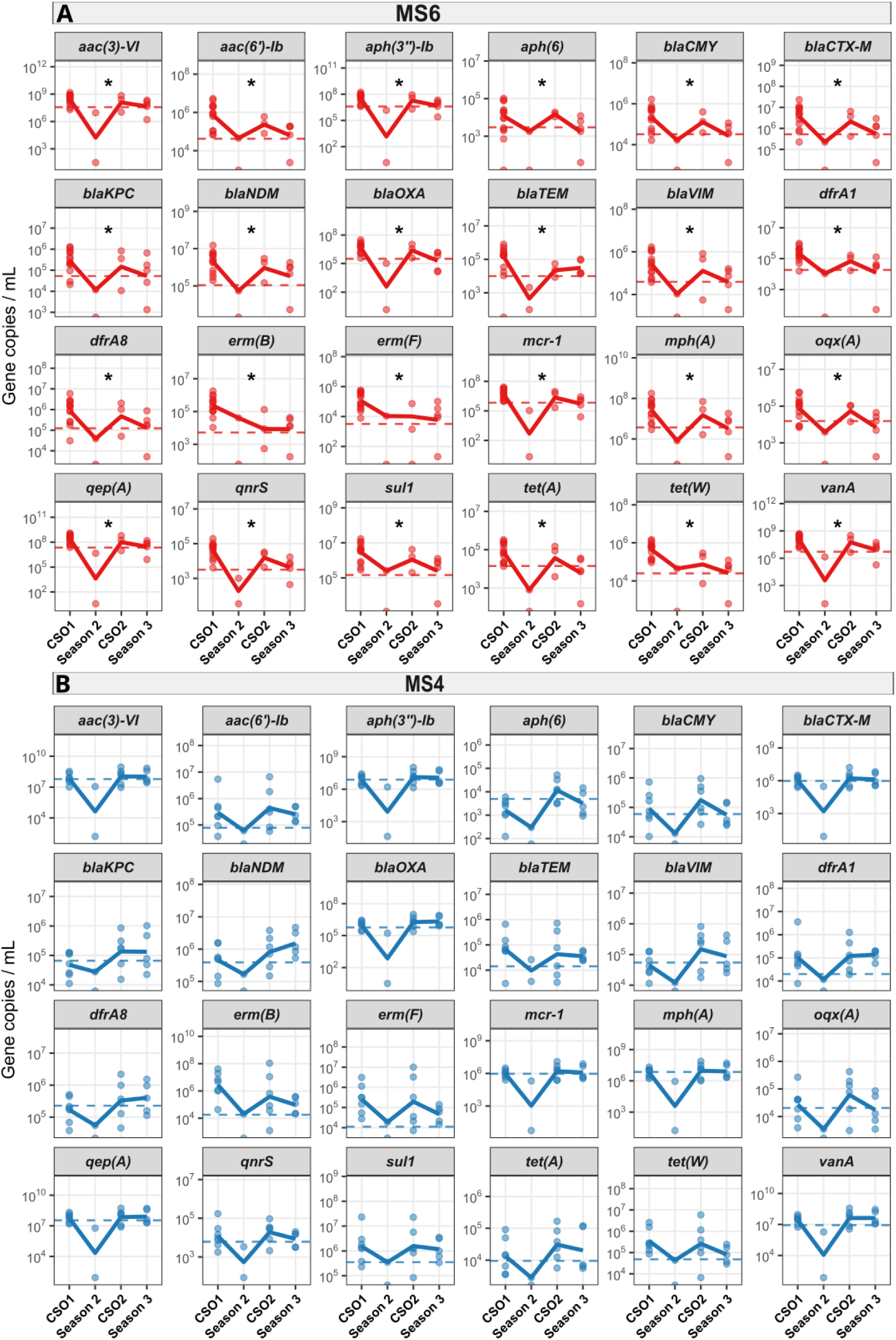
Differential ARG recovery at monitoring stations MS6 and MS4. Individual ARG abundance compared to the pre-CSO baseline (dashed line) during CSO events, and recovery phases at (A) MS6 and (B) MS4. Significant differences of phases to the baseline were determined by a Kruskal-Wallis test with Dunn’s post-hoc correction: *adjusted P<0.05, **adjusted P<0.01, ***adjusted P<0.001.

Recovery dynamics at MS4 were notably more heterogeneous, primarily because the initial impact of the overflow was highly gene-specific. None of the CSO-enriched ARGs differed significantly from CSO conditions during recovery (adjusted P>0.05 for all 24 genes), reflecting the high variance in the recovery pool driven by episodic post-event surges described above. Importantly, no ARG differed significantly from the pre-event baseline during recovery (adjusted P>0.05 for all genes; Fig. 7B), confirming that CSO-derived resistance signals were not permanently incorporated into the downstream community. Total ARG abundance during recovery (mean 4.21±6.42 × 10^8^ copies/mL, n=7) likewise did not differ significantly from either the CSO peak (P=0.391) or the baseline (P=0.265). Pathogen abundance at MS4 declined toward but did not significantly differ from CSO conditions during recovery (P=0.270; Fig. S5B). However, recovery concentrations remained significantly elevated above the pre-event baseline (P=0.031), suggesting that pathogen concentrations may not have fully returned to pre-event levels at the downstream site within the sampling window. This finding should be interpreted cautiously, as it rests on the genus-level amplicon-based pathogen metric and a limited number of recovery samples (n=7).

However, after the second CSO event, recovery initially appeared complete but was subsequently disrupted at two independent time points, characterized by renewed elevations in total bacterial and ARG abundances (Fig. 1B, 2B). These late surges reached levels comparable to CSO conditions, even though the preceding timepoints had already returned toward baseline. Overall, these results indicate that while primary CSO effects are generally transient and followed by rapid washout of most genes, downstream river sections may experience episodic, post-event surges that prolong elevated ARG and bacterial abundance beyond the initial overflow event.

## Discussion

This study provides high-resolution temporal and spatial characterisation of how CSOs shape the microbial community and resistome of an urban stream. By combining event-triggered, time-resolved sampling with seasonal baseline monitoring at sites with differing hydrological contexts, we disentangled the magnitude, spatial attenuation, and persistence of CSO-driven disturbances. Here, CSOs act as hydrologically driven pulse disturbances, introducing large loads of sewage-associated bacteria, pathogens, ARGs, and plasmid replicons into receiving waters. Importantly, the observed patterns indicate that the impact of these events is governed primarily by physical transport and dilution processes rather than by sustained ecological restructuring of the resident microbial community.

Consistent with the ecological concept of pulse disturbances, CSO events generated abrupt and pronounced surges in microbial and resistance-associated contamination. At the upstream monitoring station, bacterial concentrations during overflow approached levels characteristic of raw wastewater [59], indicating that despite the 660 m distance to the CSO discharge pipe, sewage bacteria dominated. This interpretation is further supported by the concurrent enrichment of the human wastewater markers cotinine and metoprolol, confirming that inputs of untreated domestic sewage drove the hydrological surges. Previous source-tracking has demonstrated that CSOs serve as major acute exposure pathways, capable of increasing human-associated bacterial loads by more than an order of magnitude relative to standard stormwater runoff alone [60]. Similar sharp increases in faecal indicators, ARB, and ARG abundance during overflow or bypass events have been reported in other urban systems, including Swiss rivers receiving wastewater bypass discharges [61], culture-based assessments of multidrug-resistant bacteria during CSO events [62], and urban river monitoring under CSO influence [63,64]. CSO discharges can act as substantial short-term point sources of antibiotic-resistant bacteria in receiving waters [65]. Together, these studies support the view that overflow discharges represent episodic but intense contamination events rather than minor fluctuations around baseline conditions. This massive influx is consistent with recent city-scale water quality modelling in Dresden, which estimated that CSOs currently discharge nearly 15,000 kg of organic compounds and pharmaceutical pollutants which includes antibiotics into the local catchment annually [66]. In our system, these short-lived peaks bypassed the selective barrier of wastewater treatment, sharply increasing levels of clinically relevant ARGs and human-associated pathogens during rainfall events and thus potential exposure of humans or wildlife that get into contact with the river ecosystem. Importantly, high-frequency sampling revealed that these peaks were not monotonic but comprised multiple transient maxima within single overflow episodes, indicating rapid alternation between concentrated sewage inputs and dilution phases. This complex variability is supported by previous high-frequency sampling, which revealed that ARG transport during wet-weather events is highly dynamic and rarely follows standard ’first-flush’ patterns [67].

Furthermore, the multiple transient ARG peaks we observed are a direct biological reflection of complex in sewer hydraulics in the catchment area, where varying degrees of network connectivity cause different sub-catchments to release their loads at different intervals during a single storm [10]. The scouring and mobilization of accumulated sewer sediments during peak flows likely amplify this intra-event variability. Sewer sediments act as environmental reservoirs that are capable of accumulating a distinct, highly diverse resistome, including clinically important ARGs such as *van*A and *bla*NDM-1, which are subsequently flushed into receiving waters during high-flow storm events [68]. However, here the CSO resistome and microbiome seemed to barely differ from the one found in the stream as signals of CSO events were only detected in absolute rather than relative abundance. In contrast to the relatively stable background signal resulting from continuous WWTP effluent discharge [69], CSOs created transient contamination maxima concentrating risk into short windows during and immediately after overflow rather than elevating it continuously. Thus, the primary signature of CSOs in this study was not a gradual shift in background conditions, but the repeated superimposition of high-magnitude disturbance events on an otherwise stable system.

A central mechanistic insight from our data is that the CSO-driven increase in ARG and plasmid abundance largely reflects an influx of highly concentrated sewage-associated bacteria rather than selective amplification within the river environment after the CSO ceased. All 24 quantified ARGs at MS6, and 15 of 24 at MS4, showed significant increases in absolute abundance during overflow events. Moreover, the repeated decline of ARG concentrations toward near-baseline levels even between intra-event peaks, together with the absence of significant shifts in relative abundance for most genes, indicates that these fluctuations are driven by physical hydrological transport rather than *in situ* amplification within the river environment. This indicates that, for most resistance determinants, CSOs primarily serve as mass-transfer events from the human-associated microbiome into the receiving water body. We note that the short duration of individual overflow events itself constrains the scope for in situ selection, which requires sustained selective pressure across many bacterial generations. The stability of relative abundance during these pulses is therefore consistent with the expectation that selection is unlikely to act on such timescales, and our high-resolution data confirm this empirically rather than discriminating between two equally plausible mechanisms. This aligns with large-scale metagenomic analyses demonstrating that in anthropogenically impacted environments, elevated ARG abundances are largely driven by passive dissemination of faecal pollution rather than active, on-site environmental selection [70]. Event-based studies have similarly reported substantial increases in absolute ARG loads during overflow or bypass events, while relative abundance dynamics, when examined, have mostly revealed gene-specific enrichment patterns [61,64]. In our system, broadly distributed and highly abundant markers such as *sul*1 and *tet*W maintained stable proportional representation, suggesting that their prevalence in incoming sewage closely resembles that of the background river microbiome.

Pathogen dynamics mirrored those of ARGs: although absolute abundance increased sharply and showed multiple peaks during CSO events, relative proportions remained stable, indicating a concentrated but compositionally coherent sewage community compared to the river rather than selective expansion within the river. Plasmid-associated replicons (IncP, IncW, and IncQ) followed the same pattern, while their absolute abundances surged, their relative abundances either remained stable or significantly decreased (e.g., IncQ and IncW at MS6). Together with similar observations in other CSO-impacted systems [64], this supports the interpretation that mobility potential is primarily introduced through the influx of sewage-derived bacteria rather than amplified in situ. While the overall genus-level pathogen proportion remained stable, targeted qPCR showed that the canonical faecal indicators *E. coli* and *Enterococci* were disproportionately enriched in relative terms, confirming that CSO inputs were compositionally distinct from the background stream community in terms of indicator organisms.

Building on the interpretation that CSOs primarily function as mass transfer events, the spatial contrast between MS6 and MS4 demonstrates that hydrological dilution is the dominant mechanism controlling downstream attenuation of contamination. Despite MS4 being located downstream of additional CSO outlets, both bacterial and ARG fold-enrichment were consistently lower than at MS6. This pattern is best explained by the substantially larger water volume passing through the downstream reach. MS4’s wider channel and the cumulative contribution of stormwater runoff and discharges from separate networks and streets along the 6 km stretch [71] result in greater dilution of sewage derived inputs, reducing bacterial and ARG concentrations even when total loads remain high. Similar downstream dilution has been observed in wastewater bypass and CSO-impacted systems [61,63], highlighting that physical transport processes often dominate contaminant gradients over biological filtering. This hydrological control is further reflected in the limited structural shifts observed in the resistome and overall microbial community. At MS6, CSO events produced measurable but modest changes in resistome composition, whereas at MS4, these compositional differences were no longer statistically significant. Phylum-level community structure remained largely stable at both sites. Comparable resilience of community structure despite substantial contaminant input has been observed in other CSO-influenced rivers [64], suggesting that sewage-associated bacteria introduced during overflow events largely represent a transient superimposition on the resident community rather than a competitive population capable of displacing established taxa. Crucially, the rapid return to baseline, both between intra-event peaks and after event cessation, indicates that recovery dynamics mirror the dilution-driven processes observed during contamination, further supporting the interpretation that physical transport, rather than ecological restructuring, governs system behaviour.

While the recovery at the upstream site was consistent across both events, the downstream site exhibited two isolated post-event surges in bacterial and ARG abundance following the second CSO. These elevations occurred after apparent initial recovery and in the absence of recorded upstream overflow, distinguishing them from the intra-event oscillations observed during active overflow events. Such downstream variability likely reflects additional local hydrological inputs, such as runoff or tributary-driven pulses [29], rather than delayed rebound or prolonged persistence of CSO-derived material. Several mechanisms could plausibly contribute, including the delayed arrival of a resuspended sediment plume mobilised by an earlier upstream disturbance, or differing travel times that cause individual sub-catchments to deliver their loads to the downstream reach at staggered intervals [10]. However, no rainfall, water-level, or chemical-marker data were available to corroborate a specific driver for these two timepoints, and they therefore remain unexplained within the current monitoring framework. The dominant pattern across both events nevertheless remains rapid attenuation and return toward baseline, indicating that the downstream anomaly may primarily represent episodic hydrological variability rather than sustained persistence of CSO-derived contamination.

A limitation of this study lies in the temporal and analytical scope of the monitoring design. Although high-frequency sampling resolved intra-event oscillations and short-term recovery, longer-term surveillance across multiple seasons and years would be required to assess the cumulative effects of repeated CSO exposure. In addition, our analyses relied on targeted qPCR markers and amplicon-based community profiling, which do not capture the full genomic context of resistance determinants or enable strain-level tracking of potential invaders. A further limitation is that the amplicon-based pathogen estimate is derived from genus-level 16S rRNA classification against a reference list of human-pathogenic genera. Because many of these genera also contain widespread, harmless environmental species that are common in freshwater, this metric reflects the proportion of potential human-pathogenic genera rather than confirmed pathogens; the targeted qPCR of *E. coli*, *Enterococci*, and *K. pneumoniae* provides species-level resolution that counterbalances this limitation. While the consistent return toward baseline and absence of sustained compositional shifts argue against long-term establishment of sewage-derived populations, metagenomic or cultivation-based follow-up on sediment or biofilm communities would be necessary to definitively exclude rare persistence or horizontal gene transfer events below the detection threshold of the present approach.

From a public health and management perspective, our findings indicate that CSOs represent acute but transient contamination events rather than drivers of persistent resistome restructuring under the conditions studied here. The primary risk arises from short-term exposure to high pathogen densities and clinically relevant ARGs during overflow episodes, particularly in areas proximal to discharge points. In contrast to studies reporting sustained increases in relative ARG abundance or long-term enrichment under continuous wastewater discharge or chronic disturbance scenarios, where constant effluent exposure can persistently enrich the downstream resistomes [37] and promote resistance determinants to integrate into river biofilms [72], our results show limited evidence for stable establishment or selective amplification within the river microbiome. Where persistent stressors maintain elevated relative ARG levels, invasion and long-term incorporation into environmental communities are more likely. In the present system, rapid hydrological flushing and dilution appear to limit the development of such sustained selective conditions. These findings suggest that management strategies should prioritise reducing the frequency and magnitude of overflow events to minimise acute exposure peaks, while recognising that the long-term ecological imprint of individual CSOs may be limited in hydrologically dynamic systems.

In summary, the findings indicate that CSOs act as hydrologically driven pulse disturbances, transiently increasing microbial resistance and mobility-associated loads in urban streams, but provide no evidence of fundamental restructuring of the resident microbial community. The observed intra-event oscillations, rapid downstream attenuation, and consistent return to baseline levels suggest that physical transport and dilution are the primary factors shaping system dynamics. In contrast to systems with chronic wastewater inputs, where sustained selective pressures can promote invasion and long-term ARG enrichment, the hydrodynamic conditions in this study appear to limit the stable establishment of sewage-derived resistance determinants. These results support the interpretation that CSOs primarily generate acute, short-term increases in pathogen and ARG concentrations, rather than persistent ecological transformation of the resistome. Given projections of increased extreme rainfall events, reducing the frequency and volume of overflows may be important for mitigating episodic AMR exposure in urban waterways.

## Acknowledgements

This work was supported by the Urban Resistome project funded by the Deutsche Forschungsgemeinschaft (DFG) under grant number 460816351. It further received support by the Explore-AMR project and the JPIAMR projects SEARCHER & TEXAS, funded by the German Bundesministerium für Forschung, Technologie und Raumfahrt (01KI2401, 01KI2404A & 01DO2200). Operation and maintenance of the monitoring stations were funded by the Helmholtz Water Network. Responsibility for the information and views expressed therein lies entirely with the authors. The authors thank Steffen Kunze, Christiane Zschornack, and Juliane Isler for technical support in the laboratory. The help of Stephan Becker and Alexander Bartusch during sampling is much appreciated.

## Data Availability

The datasets supporting the conclusions of this article are included within the article and its additional files or available through the corresponding author upon reasonable request. Original sequencing data is available in the NCBI sequencing read archive under project accession number PRJNA1450404.

## Author Contributions

**DK:** Conceptualization; Validation; Formal analysis; Investigation; Data Curation; Writing - Original Draft; Writing - Review & Editing; Visualization; **RPM:** Conceptualization; Methodology; Investigation; Resources; Data Curation; Writing - Review & Editing; Project administration; **SS:** Methodology; Formal analysis; Data Curation; Writing - Review & Editing; **DKn:** Investigation; Writing - Review & Editing; **FT:** Investigation; Writing - Review & Editing; **JB:** Methodology; Resources; Writing - Review & Editing; Funding acquisition; **GTV:** Validation; Writing - Review & Editing; **EDE:** Investigation; Writing - Review & Editing; **RO:** Methodology; Formal analysis; Data Curation; Writing - Review & Editing; **PK:** Conceptualization; Resources; Writing - Review & Editing; Supervision; Project administration; Funding acquisition; **TUB:** Conceptualization; Resources; Writing - Review & Editing; Supervision; Project administration; Funding acquisition; **UK:** Conceptualization; Formal analysis; Data Curation; Writing - Original Draft; Writing - Review & Editing; Visualization; Supervision; Project administration; Funding acquisition

