## Supplementary information for "Cycles of contamination and recovery: Combined sewer overflows drive acute but transient antimicrobial resistance exposure in an urban stream"

**This supplementary information contains:**

- 8 Pages
- 1 Table
- 5 Figures

**Table S1:** Primer sets used for resistomap analysis

| Gene/Marker | Target | Forward Primer | Reverse Primer |
|---|---|---|---|
| 16S rRNA | 16S rRNA | GGGTTGCGCTCGTTGC | ATGGYTGTCGTCAGCTCGTG |
| *aac(6')*-Ib | Aminoglycoside | GTTTGAGAGGCAAGGTACCGTAA | GAATGCCTGGCGTGTTTGA |
| *aac3*-VI | Aminoglycoside | CGTCACTTATTCGATGCCCTTAC | GTCGGGCGCGGCATA |
| *aph3*-ib | Aminoglycoside | AACAGGTTTGGGAGGCGATG | CGCAACAAGCCTCTCCTGAA |
| *aph*6 | Aminoglycoside | CCCATCCCATGTGTAAGGAAA | GCCACCGCTTCTGCTGTAC |
| $bla_{CMY}$ | Beta Lactam | AAAGCCTCAT GGGTGCATAAA | ATAGCTTTTGTTTGCCAGCATCA |
| $bla_{CTX\text{-}M}$ | Beta Lactam | CGTACCGAGCCGACGTTAA | CAACCCAGGAAGCAGGCA |
| $bla_{KPC}$ | Beta Lactam | GCCGCCGTGCAATACAGT | GCCGCCCAACTCCTTCA |
| $bla_{NDM}$ | Beta Lactam | GGCCACACCAGTGACAATATCA | CAGGCAGCCACCAAAAGC |
| $bla_{OXA}$ | Beta Lactam | TGTTTTTGGTGGCATCGAT | GTAAMRATGCTTGGTTCGC |
| $bla_{TEM}$ | Beta Lactam | CGCCGCATACACTATTCTCAG | GCTTCATTCAGCTCCGGTTC |
| $bla_{VIM}$ | Beta Lactam | GCACTTCTCGCGGAGATTG | CGACGGTGATGCGTACGTT |
| *dfr*A1 | Trimethoprim | GGAATGGCCCTGATATTCCA | AGTCTTGCGTCCAACCAACAG |
| *dfr*A8 | Trimethoprim | GGTCGCACCTGCATCGTTA | AGCGCCACCAATGACGTAG |

| | | | |
|---|---|---|---|
| *erm*B | MLSB | TAAAGGGCATTTAACGACGAAACT | TTTATACCTCTGTTTGTTAGGGAATTGAA |
| *erm*F | MLSB | CAGCTTTGGTTGAACATTTACGAA | AAATTCCTAAAATCACAACCGACAA |
| *mcr*1 | Other | CACATCGACGGCGTATTCTG | CAACGAGCATACCGACATCG |
| *mph*A | MLSB | CTGACGCGCTCCGTGTT | GGTGGTGCATGGCGATCT |
| *oqx*A | MDR | GAGTCAACCTACCTCCACTATCA | GCTGCGAGTTATCCAGCAG |
| *qep*A | Quinolone | GGGCATCGCGCTGTTC | GCGCATCGGTGAAGCC |
| *qnr*S | Quinolone | CCACTTTGATGTCGCAGATCTTC | CCCTCTCCATATTGGCATAGGAAA |
| *sul*1 | Sulfonamide | GCCGATGAGATCAGACGTATTG | CGCATAGCGCTGGGTTTC |
| *tet*A | Tetracycline | GCTGTTTGTTCTGCCGGAAA | GGTTAAGTTCCTTGAACGCAAACT |
| *tet*W | Tetracycline | ATGAACATTCCCACCGTTATCTTT | ATATCGGCGGAGAGCTTATCC |
| *van*A | Vancomycin | GGGCTGTGAGGTCGGTTG | TTCAGTACAATGCGGCCGTTA |
| IncP_oriT | MGE | CAGCCTCGCAGAGCAGGAT | CAGCCGGGCAGGATAGGTGAAGT |
| IncQ_oriT | MGE | TTCGCGCTCGTTGTTCTTCGAGC | GCCGTTAGGCCAGTTTCTCG |
| IncW_trwAB | MGE | AGCGTATGAAGCCCGTGAAGGG | AAAGATAAGCGGCAGGACAATAACG |

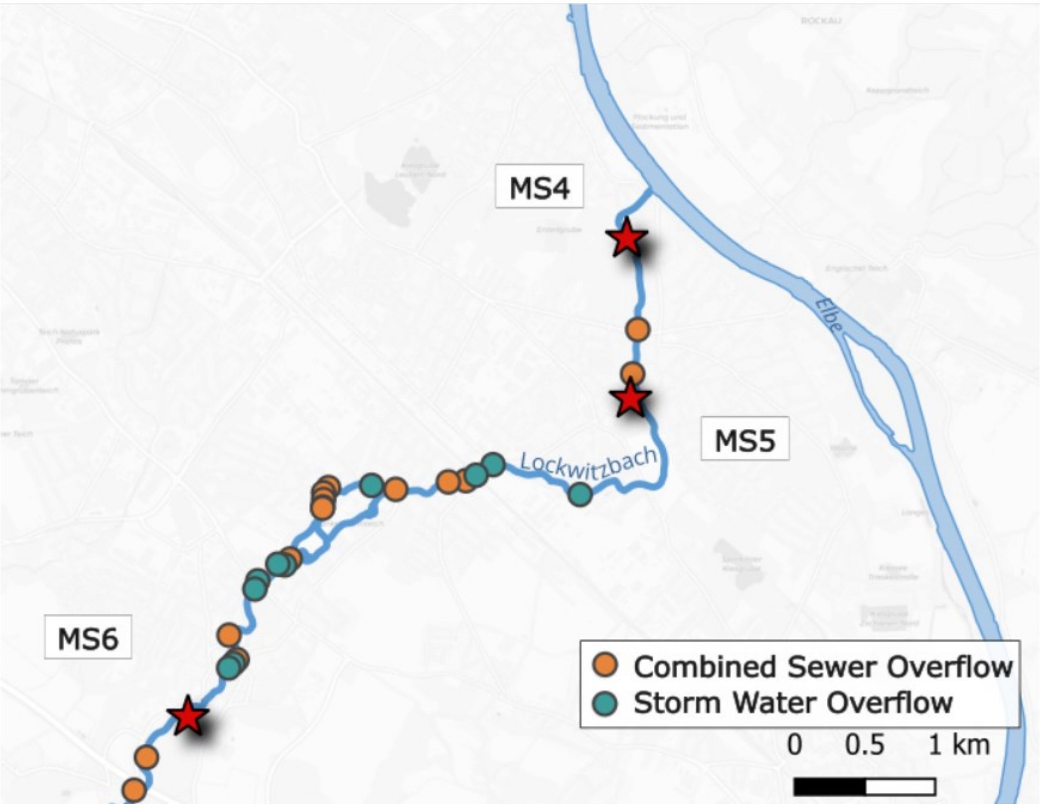


**SI Figure 1:** Location of the three monitoring stations along the Lockwitzbach, a 24 km urban tributary of the Elbe River in Dresden, Germany. MS6 (upstream) and MS4 (~6 km downstream, near the Elbe confluence) are CSO-impacted stream stations. MS5 (~4.5 km downstream of MS6) is a stormwater-only (SWO) site on a separate storm-sewer network that receives only urban surface runoff. Other stormwater discharge points are shown as green circles and other CSO discharge points as orange circles.

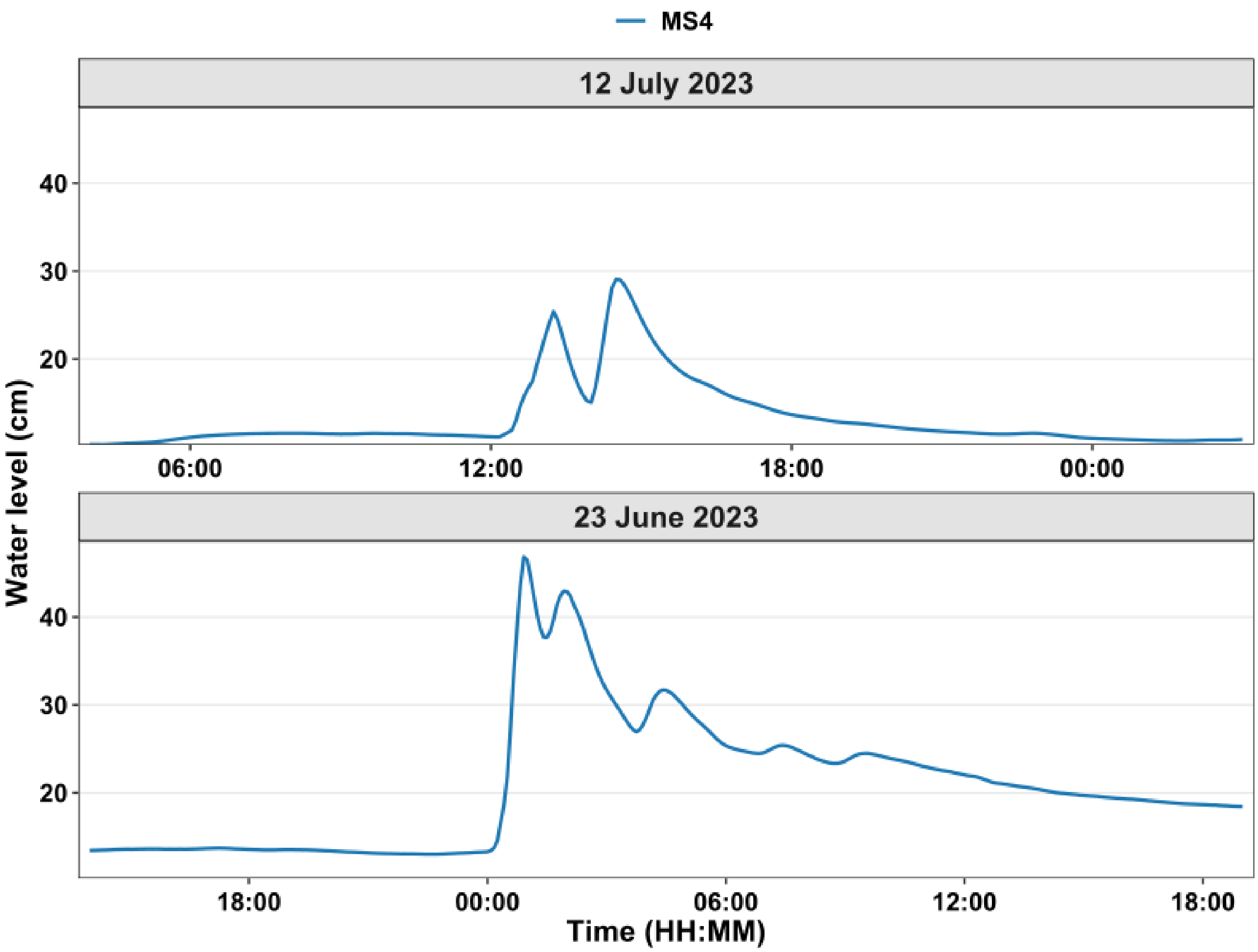


**SI Figure 2:** Water level dynamics during CSO events. Water Level (cm) recorded at 5-minute intervals at MS4 for the two events analyzed in this study.

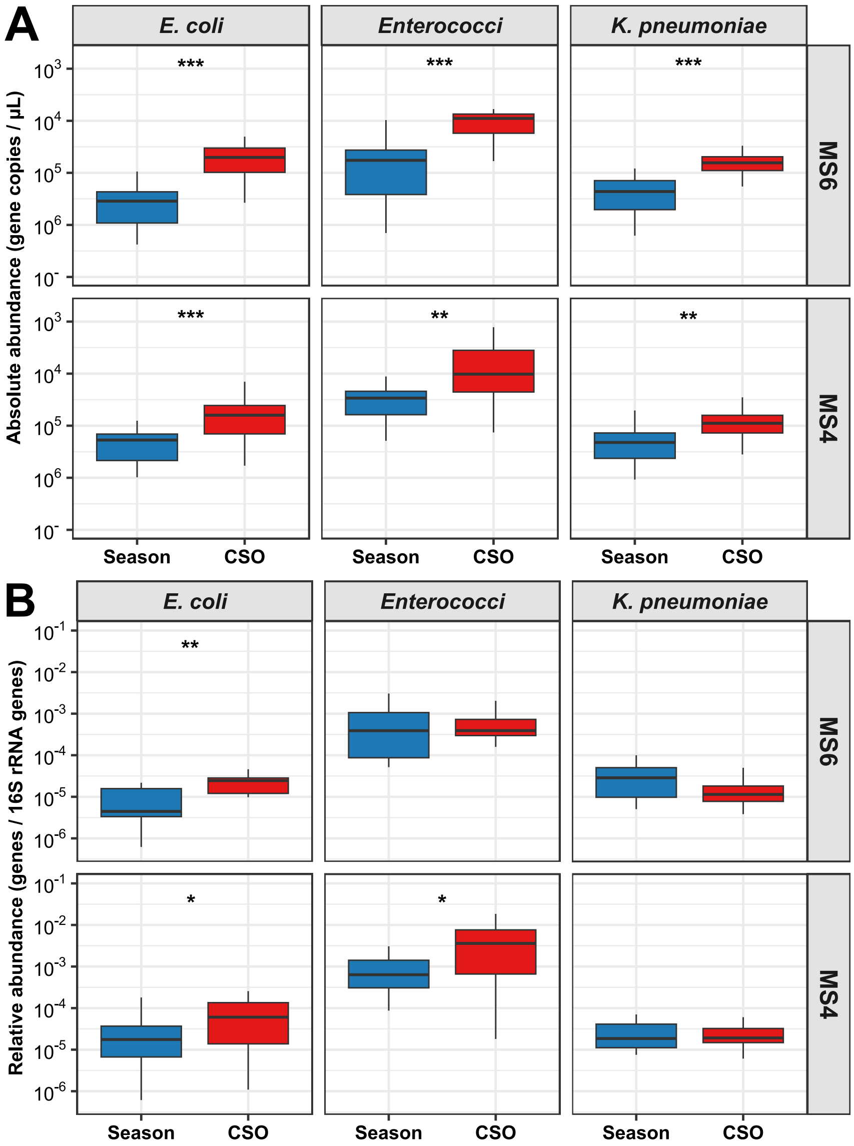


**SI Figure 3:** (A) Absolute abundance (gene copies/µL) and (B) Relative abundance, normalised to the 16S rRNA gene of E. coli, Enterococci, and Klebsiella pneumoniae. Adjusted P-values from Mann–Whitney U tests with Benjamini–Hochberg FDR correction are indicated above each comparison (*adjusted P<0.05, **adjusted P<0.01, ***adjusted P<0.001).

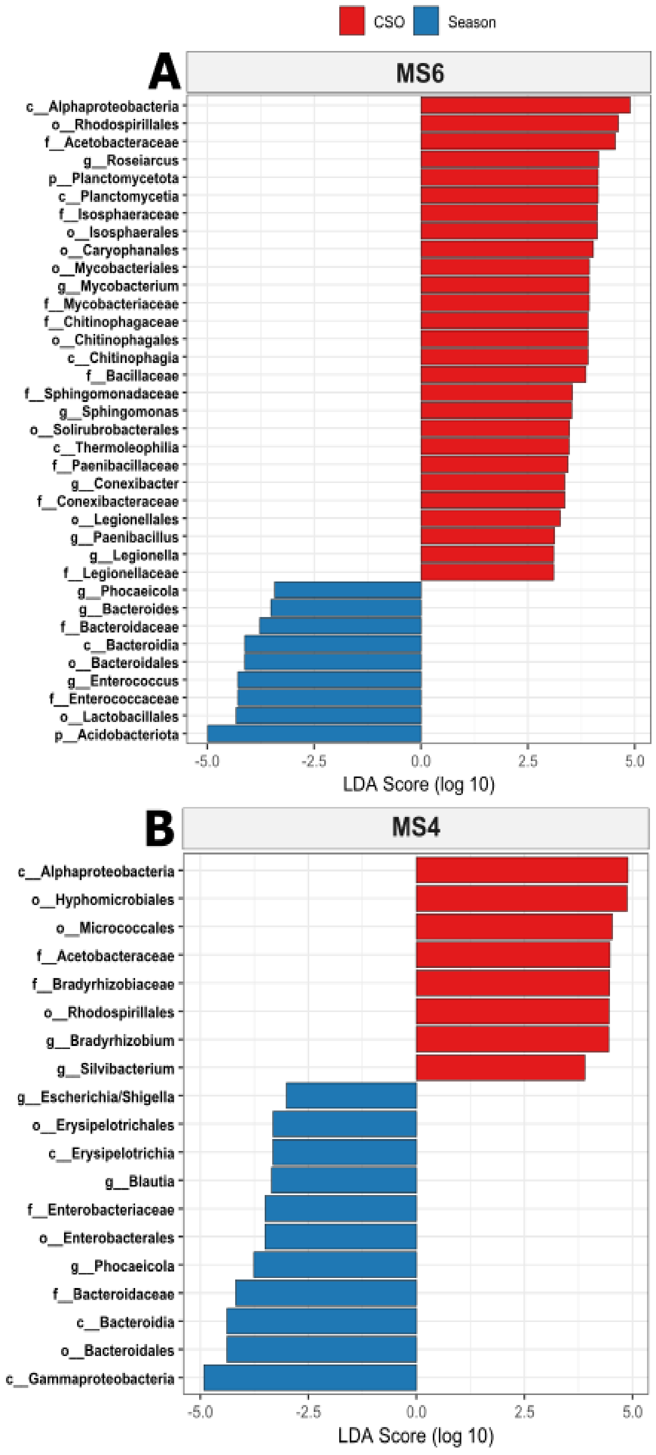


**SI Figure 4:** Linear Discriminant Analysis Effect Size (LEfSe) showing differentially abundant bacterial taxa distinguishing CSO events from seasonal conditions at MS6 and MS4. Horizontal bars represent the effect size for taxa meeting the significance threshold (LDA>3.0, P<0.05). Colours indicate the condition in which the taxon was significantly enriched (Red=CSO-associated; Blue=season-associated).

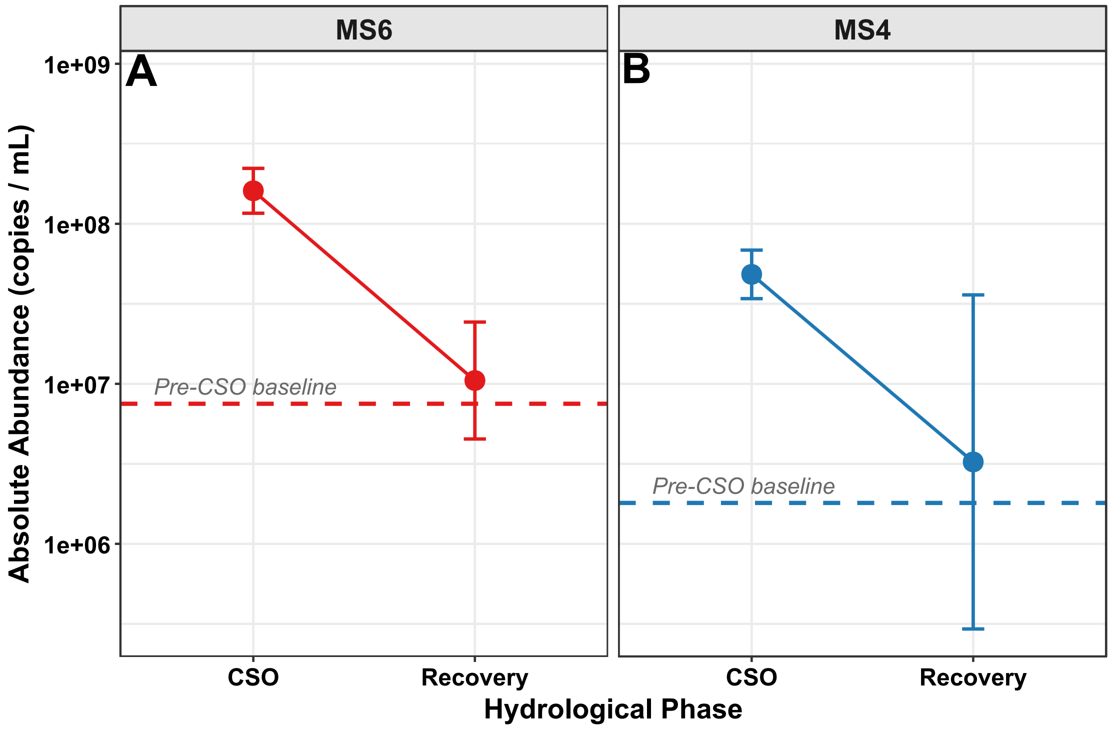


**SI Figure 5:** Pathogen abundance during recovery. Absolute pathogen abundance (copies/mL) during pre-CSO baseline, CSO event, and recovery phases at (A) MS6 and (B) MS4.
